# Adaptations of gram-negative and gram-positive probiotic bacteria in engineered living materials

**DOI:** 10.1101/2025.02.19.638224

**Authors:** Varun Sai Tadimarri, Tanya Amit Tyagi, Cao Nguyen Duong, Sari Rasheed, Rolf Müller, Shrikrishnan Sankaran

## Abstract

Encapsulation of microbes in natural or synthetic matrices is a key aspect of engineered living materials, although the influence of such confinement on microbial behavior is poorly understood. A few recent studies have shown that spatial confinement and mechanical properties of the encapsulating material significantly influence microbial behavior, including growth, metabolism, and gene expression. While such effects have been shown to elicit various responses in a few micro-organisms like *E. coli*, yeast, and cyanobacteria, systematic comparative studies between different organisms in the same confinement conditions are missing. Thus, in this study, we report the adaptive responses exhibited by rod-shaped gram-negative and gram-positive probiotic bacteria that are of great interest for developing therapeutic engineered living materials. Accordingly, gram-negative *E. coli* Nissle 1917 and gram-positive *L. plantarum* WCFS1 were encapsulated in hydrogel matrices and their growth, metabolic activity, and recombinant gene expression were investigated. By varying the polymer concentration and degree of chemical cross-linking in the hydrogels, it was possible to modulate their stiffness and study how the bacteria adapted to these different confinement conditions. In accordance with previous reports, both bacteria grow from single cells into confined colonies but more interestingly, in *E. coli* gels, mechanical properties influenced colony growth, size, and morphology, whereas this did not occur in *L. plantarum* gels. However, with both bacteria, increased matrix stiffness led to higher levels of recombinant protein production within the colonies. By measuring metabolic heat generated in the bacterial gels using a novel isothermal microcalorimetry technique, it was inferred that *E. coli* adapts to the mechanical restrictions through multiple metabolic transitions and is significantly affected by the different hydrogel properties. Contrastingly, both these aspects were not observed with *L. plantarum*. These results revealed that despite both these bacteria being gut-adapted probiotics with similar geometries, mechanical confinement affects them considerably differently. The weaker influence of matrix stiffness on *L. plantarum* is attributed to its slower growth and thicker cell wall possibly enabling the generation of higher turgor pressures to overcome restrictive forces under confinement. By providing fundamental insights into the interplay between mechanical forces and bacterial physiology, this work advances our understanding of how matrix properties shape bacterial behavior. The implications of these findings will aid the design of engineered living materials for therapeutic applications.

## Introduction

The interdisciplinary field of Engineered Living Materials (ELMs) has experienced significant growth in recent years, combining concepts and techniques from synthetic biology and material science. Several proof-of-concept studies have displayed the adaptability of ELMs, demonstrating their broad-spectrum applicability in diverse sectors, from construction to biomedicine. ^**[1] [2]**^ A common feature shared by several ELMs is the mechanical confinement of microorganisms within materials. Recent studies have shown that encapsulating microbes in hydrogels can affect their growth and metabolism. This confinement-induced alteration of microbial behavior may influence the overall performance and functionality of the ELM. ^**[3-9]**^ Common observations of these studies are that microbes within hydrogels grow as dense colonies that may or may not be restricted in size depending on the mechanical properties of the surrounding matrix and availability of nutrients. Moreover, as the depth of the hydrogel increases, the diffusion of essential nutrients and gases (such as oxygen and carbon dioxide) becomes limited/restricted. Such limitations in resources lead to a variation in bacterial colony size, where microorganisms situated deeper within the hydrogel matrix tend to be smaller in size compared to those closer to the surface. ^**[4] [9] [10] [11]**^ Furthermore, mechanical confinement has been shown to enhance the expression of genes associated with the antibiotic resistance,^**[10]**^ and maintain particular phenotypic states in microbes like yeast and cyanobacteria. ^**[4] [12]**^ Our previous studies have shown that encapsulating an *Escherichia coli* strain (ClearColi) in Pluronic F127 diacrylate (PluDA) hydrogels affected the growth rate, morphology, and protein expression of bacteria. More importantly, by modulating the degree of covalent cross-linking in these hydrogels, we observed that bacterial parameters were directly influenced by the network’s mechanical properties. Increasing degrees of covalent cross-linking led to a decrease in growth rate and colony size, and an increase in colony sphericity. Surprisingly, protein production rates exhibited a non-monotonic response, peaking at intermediate degrees of covalent cross-linking. This suggested that under these conditions, growth was sufficiently slowed to metabolically favor protein production. At the highest degree of covalent cross-linking, lower production rates result from bacteria allocating resources to overcome constrictive mechanical forces. These findings highlighted the need to optimize mechanical properties of the encapsulating material for optimal performance of an ELM. Additionally, these studies unsurprisingly reveal that different microbes show differential responses to confinement. However, since each study involves a single microbial species and different hydrogel components, it cannot be determined how the same mechanical confinement conditions will influence different microbes. While some studies have investigated co-culture ELMs with different microbial species encapsulated in hydrogels and performing symbiotic functions, these findings have not compared the influence of confinement on the individual species. ^**[13-16]**^ Investigating the effects of mechanical confinement on different microbial species will enable us to uncover species-specific adaptations and interactions within ELMs and aid in finding optimal material-microbe combinations for different applications.

In this perspective, this study compares the effect of mechanical confinement on two distinct bacterial species: Gram-negative *E. coli* ^**[17]**^ and the Gram-positive *Lactiplantibacillus plantarum* ^**[18] [19]**^. Both organisms are rod-shaped and exhibit the ability to survive and grow in both aerobic and anaerobic environments. *E. coli* is the predominant bacterial species utilized in ELM research, while *L. plantarum* is a widely studied probiotic strain with beneficial properties for humans, animals, and plants. ^**[20-22]**^ *L. plantarum* also has significant industrial use in food fermentation and lactic acid production. ^**[23] [24]**^ In this study, we specifically chose the probiotic strains, *E. coli* Nissle 1917 and *L. plantarum* WCFS1, which are genetically tractable and are of considerable interest for the development of ELMs towards biomedical applications. Despite their similarities, the strains exhibit significant differences in their cellular architecture and physiology, including, cell-wall composition, cell membranes, cytoplasmic turgor pressure, and metabolic pathways. ^**[25-27]**^ Given these distinct cellular characteristics, this study aimed to explore how *E. coli* Nissle 1917 and *L. plantarum* WCFS1 respond differently to mechanical confinement. To create the different mechanical conditions, we employed PluDA hydrogels, expanding upon our previous research by varying both the degree of chemical cross-linking and polymer concentration, resulting in nine distinct experimental conditions. The bacterial responses were assessed in terms of colony growth, morphology, and protein production (confocal microscopy) and in terms of metabolic activity (isothermal microcalorimetry). These analyses suggested that probiotic *E. coli* and *L. plantarum* share some common responses to mechanical confinement but are likely to employ distinct mechanisms to grow and maintain activity under spatial constraints.

## Experimental Section

### Preparation of precursor solutions

Pluronic diacrylate (PluDA) was synthesized by reaction of Pluronic F127 (Plu) with acryloyl chloride in the presence of triethylamine according to a previously reported protocol ^**[39]**^ to achieve a functionalization degree of 90%. 23% (w/w) Plu and PluDA stock solutions were prepared separately in two different nutrient broths, LB medium (for *Escherichia coli Nissle* 1917 experiments) and MRS medium (for *Lactiplantibacillus plantarum* WCFS1 experiments). All the solutions were supplemented with Irgacure 2959 photo initiator (0.2% w/v) to facilitate the cross linking when exposed to UV radiation. 16.7% and 20% solutions of Plu and PluDA were prepared by diluting 23% stock solutions with respective solvent (LB or MRS medium). All the solutions were stored at 4 °C until use.

### Rheology of polymer combinations

A rotational rheometer (DHR3, TA instruments) was used to measure the rheological properties. A 20 mm peltier plate (Serial number 106155, transparent to UV) was used as a bottom plate and 12 mm stainless steel disk was used as top plate. Experiments were performed at the room temperature (25 °C), and a UV source (Omnicure, Series 1500, 365 nm, 6 mW/cm^2^) was equipped with rheometer. A gap of 300 µm between the plates and a volume of precursor polymer solution of 35 µL was used for the experiment. After the sample was loaded, 2 minutes of UV exposure was programmed, followed by test measurements after 10 minutes of sample loading. Strain sweeps were conducted from 0.1% to 1000% at a frequency of 1 Hz. From these results, linear viscoelastic region was identified and storage modulus per polymer condition was plotted using data obtained from the plots.

### Genetic modification of bacterial strains and culturing the bacteria

*E. coli* Nissle 1917 and *L. plantarum* WCFS1 strains were engineered with p256-mCherry plasmids to obtain the fluorescent constructs of respective strains. *E*.*coli Nissle* was transformed with p256-mCherry with kanamycin selection marker and *L. plantarum* wcfs1 was transformed with p256-mCherry with erythromycin selection marker. Competent cells of both strains and transformation were performed using the published protocols. ^**[35][36]**^ Respective colonies of both strains were inoculated either from agar plate or from their glycerol stocks. *E*.*coli Nissle* 1917 was grown in LB medium supplemented with kanamycin antibiotic (50 µg/mL) whereas *L. plantarum* was grown in MRS medium supplemented with erythromycin (10 µg/mL) for 14 - 16 hours at 37 °C followed by subculturing the bacteria on the next day to OD 0.1 and let it grow till log phase (0.5-0.8) before encapsulation.

### Microscopy of bacterial culture

Grown cultures of *E. coli* Nissle 1917 and *L*. plantarum WCFS1 strains expressing mCherry were diluted to OD 1 and 200 µL of diluted cultures were pipetted into a 96-well black plate with transparent bottom. Fluorescence microscopy of these bacterial cultures was performed using a table-top Keyence microscope (BZ-X800E). Multichannel images were captured at 100x magnification using the CH3 (red fluorescence) and CH4 (brightfield) channels, the images were analysed using imageJ.

### Bacterial encapsulation for microscopy

Overnight incubated cultures of *E. coli* Nissle 1917 and *L. plantarum* WCFS1, both expressing mCherry fluorescent protein were stopped at optical density (OD_600_) of 0.6-1 and suspended in specific composition of polymer precursor solutions (at 4 °C) in the volume ratio of 9:1 (Polymer solution: Bacterial culture) to achieve a final OD_600_ of 0.01 (*E. coli* Nissle 1917: 1*10^4^ CFU; *L. plantarum* WCFS1: 6*10^3^ CFU) within the gels having final (w/w) polymer concentrations of 15%, 18%, and 21%. This mix was then vortexed immediately before placing on ice to ensure that the pluronic solutions remain in their liquid state. 10 µL of polymer-bacteria suspension was pipetted into the well of a Ibidi 96 Well µ-Plate. Once all wells were pipetted with respective samples, the plate was transferred to UV chamber and exposed to UV radiation (365 nm) for 2 minutes at an intensity of 6 µW/cm^2^. 30 µL of silicone oil (350 cSt, Sigma-Aldrich) was pipetted in all wells with polymerized gels to prevent the drying of the hydrogel during the experiment. The plate was incubated at 37 °C until it was transferred to microscopy analysis.

### Kinetic microplate reader measurements to study bacterial growth and recombinant protein production rates

Engineered *E. coli* Nissle 1917 and *L. plantarum* WCFS1 were encapsulated in different Plu/PluDA formulations as mentioned in the section above. Crosslinked bacterial gels of 10 µL volume were fabricated in an Ibidi 96 Well µ-Plate, covered by 30 µL of silicone oil. The plate was covered, and an overnight growth kinetics were set for a duration of 24 hours (one measurement every hour) to measure the OD600 and fluorescence intensity of mCherry expression (Ex/Em: 587/625 nm) from the bacterial hydrogels with a gain of 100 and Z-position of 21760 µm. The obtained raw data was analysed using GraphPad prism to plot the growth rate and fluorescence intensity plots.

*Growth rate = (OD600 at T2 – OD600 at T1)/ ΔT*

*Fluorescence intensity rate = (Fluorescence units at T2 – Fluorescence units at T1)/ ΔT*

### Image acquisition and data analysis for colony volume, colony sphericity and fluorescence intensity

Samples were imaged using a Zeiss Cell Discoverer 7 microscope equipped with a ZEISS LSM 900 and AiryScan 2 (Zeiss, Oberkochen, Germany). Fast AiryScan mode was used with a Zeiss Plan-Apochromat 20×/0.95 objective and a 1x tube lens (Optovar). Excitation and emission wavelengths were set to 561 nm and 565–700 nm, respectively, to image mCherry-expressing bacterial populations. Each image covered an area of 197.91 × 197.91 µm (1156 × 1156 pixels) in the XY plane, with a 100 µm Z-stack acquired using ZEN 3.6 Blue software (Zeiss). Colony volume, sphericity, and fluorescence intensity were analyzed using the Imaris Surface module with automatic thresholding (v10.1, Bitplane, Zurich, Switzerland).

The data was obtained through IMARIS Imaging software and went through a series of steps to extract meaningful insights. The data processing and analysis were carried out employing certain Python libraries. Initially, the dataset was loaded from a set of CSV files using the Pandas library. ^**[30]**^ Specific parameters such as Volume, Sphericity, and Intensity mean were extracted into separate DataFrames for each condition. Only bacterial colony volumes greater than 10 µm^3^ were selected for analysis, ensuring that the focus remained on the most significant data points. After this initial data selection, a bootstrapping technique, utilizing NumPy,^**[31]**^ was utilized to create a distribution of resampled values for each condition. With these bootstrapped values, confidence intervals were formed to estimate the plausible range of values within a given level of confidence (95%). Plots were created using python libraries Matplotlib ^**[32]**^ and Seaborn ^**[33]**^ to compare the original data with the bootstrapped confidence intervals, illustrating the underlying patterns.

### Bacterial encapsulation for CalScreener based isothermal microcalorimetric analysis

These experiments were performed using the calScreener Symcel isothermal microcalorimetry instrument (Catalog #1200001). For hydrogel preparation, 50 µL of polymer-bacteria suspensions were pipetted into individual wells of a sterile 48-well calPlate™ containing single-use sterile plastic inserts (SymCel, Sweden). The plate-containing inserts were then transferred to UV chamber and exposed to UV radiation (365 nm) for 2 minutes at an intensity of 6 µW/cm2. After cross linking, the inserts with bacterial hydrogels were loaded into titanium vials and well closed to ensure optimal performance during high-precision calorimetric testing. After device calibration, calorimetric measurements were initiated as described in literature.^**[28]**^ The data was analyzed using calView™ 2.0 software. Heat flow data from the CalScreener were evaluated based on key metrics such as maximum magnitude, peak timing, detection time, and area under the curve (AUC). These metrics were derived using Simpson’s rule with the assistance of the SciPy ^**[29]**^ and Matplotlib libraries.

### Statistical Analysis

In Figure 4, the Mann-Whitney U test was used to assess pairwise differences between nine different formulations for each of the three parameters: volume, sphericity, and intensity mean. This non-parametric test was chosen because it compares ranks without assuming normality. Our data was found to be non-normally distributed upon visualization, which justified the use of non-parametric tests. Heatmaps of the pairwise p-values between nine different formulations for sphericity, volume, and intensity mean are provided in the supplementary material **(Fig. S2, S3)**. The heatmaps were generated using the Seaborn ^**[33]**^ and Matplotlib ^**[32]**^ libraries in Python. In Figure 5, the Kruskal-Wallis Test followed by post-hoc Mann Whitney U test was performed to determine significant differences between the different conditions by testing null hypothesis. The test was performed using the following online tool - https://www.statskingdom.com/kruskal-wallis-calculator.html.

## Results and Discussion

The hydrogels for bacterial encapsulation were made of Pluronic F-127 (Plu) and its diacrylated derivative (PluDA), mixed in three different ratios (1:0 = DA0, 1:1 = DA50, 0:1 = DA100) at three different total polymer concentrations (15, 18, 21 w/w%) to yield nine different formulations (Fig. 1). Pluronic F-127 hydrogels undergo physical assembly at room temperature and the diacrylated variants can be chemically cross-linked to increase gel stability and stiffness. Thus, the mechanical properties of these hydrogels are dictated either purely by physical cross-linking (DA0), by physical & chemical cross-linking (DA50), or predominantly by chemical cross-linking (DA100). Notably, their stiffnesses ranged from a storage modulus of 3 kPa to 110 kPa (Fig. 1b).

**Figure 1:**
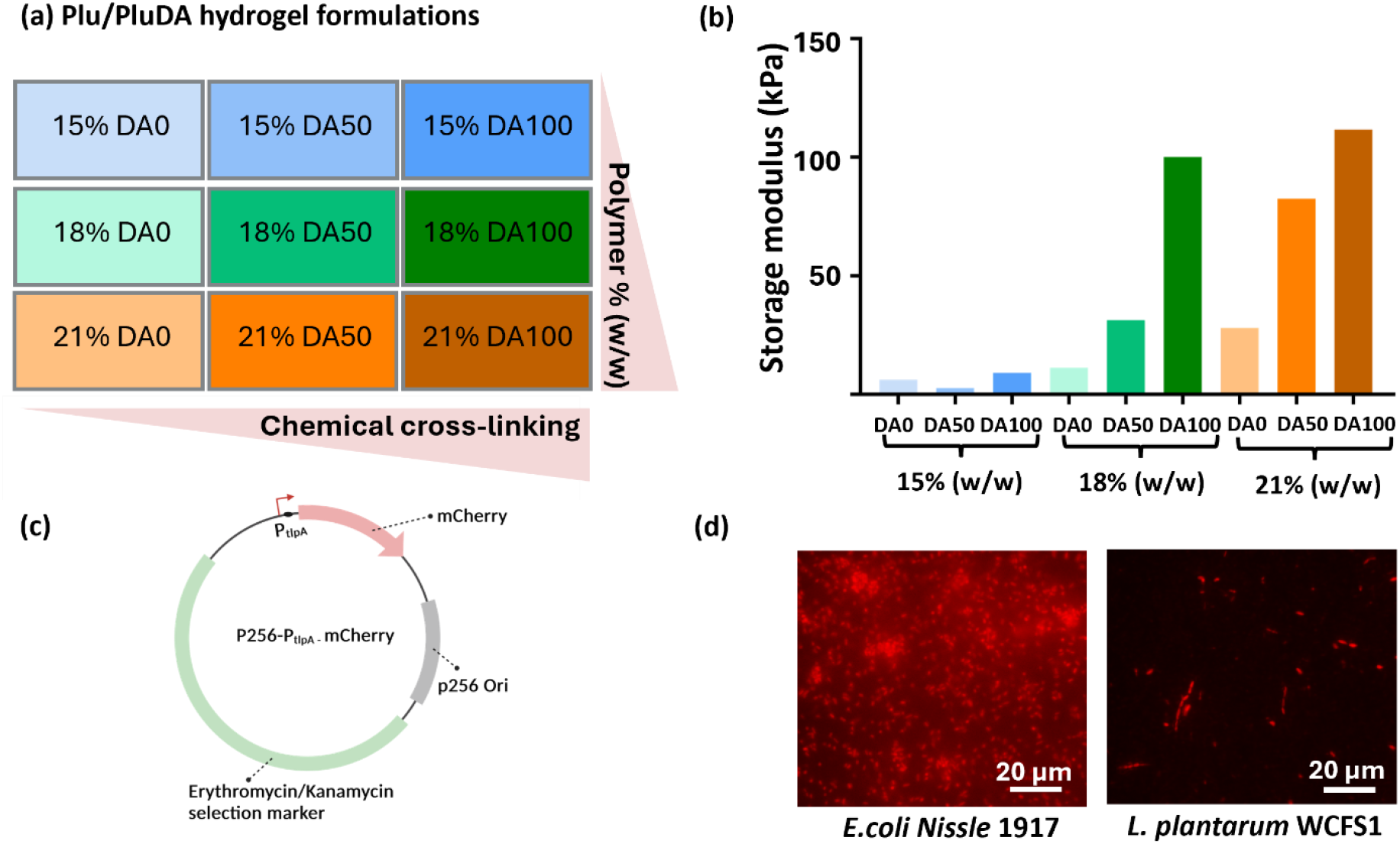
**(a)** Scheme representing the different combinations of Plu/PluDA in the hydrogels along with the colors used to represent these combinations throughout the study. DA 0 refers to 100% Plu, DA 50 refers to 50% Plu & 50% PluDA, and DA 100 refers to 100% Plu. **(b)** Bar graph representing storage modulus of the nine combinations of Plu/PluDA hydrogels in the increasing order of polymer concentration and chemical crosslinking. **(c)** Plasmid map for the plasmids with p256 origin of replication, P_tlpA_ promoter driving the expression mCherry fluorescent protein, and selection marker being either kanamycin for E.coli or erythromycin for L. plantarum. **(d)** Fluorescence microscopic images of E.coli Nissle 1917 and L. plantarum WCFS1 expressing mCherry in liquid culture captured with same microscopy settings.

The bacteria used in this study are fluorescent strains of *E*.*coli Nissle* 1917 and *L. plantarum* WCFS1 that were genetically engineered with the same plasmid coding for mCherry expression (Figure 1). This plasmid has a broad range replicon (p256) that is medium copy in *E*.*coli* (50-100) and low copy (3-5) in *L. plantarum*. The expression of mCherry is driven by the P_tlpA_ promoter with high strength in *E. coli* ^**[35]**^ and moderate strength in *L. plantarum*. ^**[36]**^ The bacteria were mixed with the polymer solutions in a volume ratio of 1:9 at 4 °C to achieve a final bacterial density of OD_600nm_ 0.01. For photo-polymerization, the hydrogels were subjected to 2 min of exposure to 365 nm that has previously been shown to negligibly affect bacterial behavior. ^**[9]**^

### 1) Growth and protein production of the bacterial population in confinement

Bacterial growth and protein production in confinement are important functions in ELM that confer them with unique capabilities. The bacterial hydrogels were prepared with the polymers dissolved in the optimized media for each species (LB medium for *E. coli* and MRS for *L. plantarum*). For high-throughput analysis, the bacterial hydrogels were formed in 96-well µ-plates 3D (Ibidi) within the lower well (10 µL) and covered with silicone oil to prevent evaporation of the water in the hydrogel (Fig. 2a). Growth kinetics were measured for 24 hours after encapsulation using a microplate reader. The first remarkable observation is that the overall growth profile was different for both bacterial strains. *E*.*coli Nissle* 1917 starts with a short lag phase for 2-4 hours followed by an expansion phase for the next 6-10 hours and then a saturation phase is reached around 12 hours, most likely due to exhaustion of the available nutrients (Fig. 2b). On the other hand, *L. plantarum* seemed to grow slower with a long lag phase for the first ten hours followed by two expansion phases with the second having a slower rate (<0.002 ΔOD_600nm_/h) than the first (0.002 – 0.004 ΔOD_600nm_/h) **(Fig. 2d)**. In comparison, when both strains were grown in well plates as non-encapsulated cultures, they exhibited a nearly negligible lag phase (<1 h), followed by two expansion phases (1-4 & 4-7 or 4-12 h) and a saturation phase (>7 or 12h) **(Fig. S1a, S1b)**. Notably, each phase was considerably shorter than what was observed with the encapsulated bacteria, indicating that mechanical confinement within the hydrogels slows down bacterial growth. In the case of encapsulated *E. coli*, the sharp transition between the expansion and saturation phases could be a result of the bacteria growing as spatially confined colonies that exhaust the nutrients surrounding them and face diffusion limitations to support the entire colony. In the case of encapsulated *L. plantarum*, such sharp transitions between the two expansion phases are discernable in certain hydrogel formulations although less prominent possibly due to their slower and extended growth phases.

**Figure 2:**
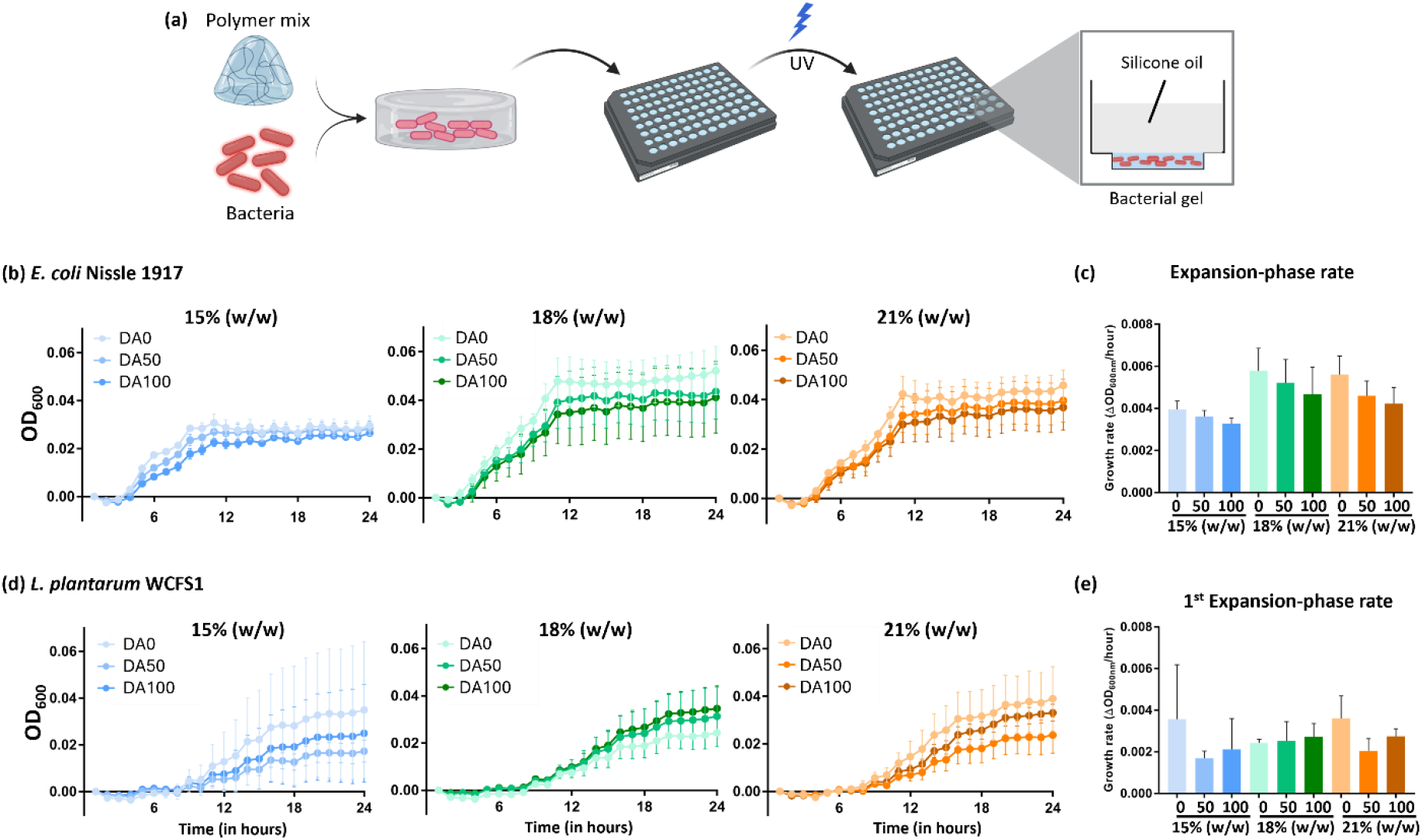
**(a)** Schematic representation of manual fabrication of bacterial hydrogels. Bacteria and polymer were mixed in a ratio of 1:10 followed by pipetting into the lower well of a 96-well well µ-plates 3D (Ibidi) and photo-crosslinking of the bacterial gels. The zoomed-in section depicts the transverse section of a well with hydrogel (10 µL) in the bottom chamber covered with 50 µL of silicone oil. **(b, d)** Normalized growth kinetics of E.coli Nissle 1917 (b) and L. plantarum WCFS1 (d) expressing mCherry in the nine Plu/PluDA combinations for 24 hours (N=4). **(c**,**e)** Column plots representing the expansion phase rate (change in OD_600nm_ per hour) of E.coli Nissle 1917 (c) and L. plantarum WCFS1 (e) across 9 polymer combinations between 4 - 9 hours of growth for E. coli and 8 - 14 hours of growth for L. plantarum (N=4).

Within each polymer concentration, total growth and rate of the expansion phase did not vary significantly from DA0 to DA100, although for *E. coli*, both parameters on average seemed to reduce with increasing degrees of chemical cross-linking (Fig. 2c), while *L. plantarum* did not exhibit any apparent trend (Fig. 2d). This could indicate that the mechanical differences between the different hydrogels possibly influence *E. coli*’s growth but not that of *L. plantarum*. A contributing factor to such a difference may be the higher turgor pressures in gram-positive bacteria (>1000 kPa) compared to gram-negative bacteria (30 – 300 kPa), ^**[27]**^ since it is the primary force driving bacterial cell expansion.

Apart from monitoring bacterial growth, the production of mCherry protein from the plasmid was quantified by measuring fluorescence intensity. Interestingly, protein production profiles were more aligned for both strains, with sequentially slow, rapid and again slow production rates, although total production was an order of magnitude higher in *E. coli* compared to *L. plantarum*. In *E. coli*, start of the rapid production phase (∼6 h) corresponded to the mid-log phase of growth but end of the rapid phase (10-12 h) nearly correlated with the saturation phase of growth **(Fig. 3a)**. Contrastingly, in *L. plantarum*, the start of the rapid production phase (∼9 h) roughly correlated with the start of the first expansion phase but the end of the first expansion phase (∼16 h) occurs a little beyond the mid rapid phase of production (Fig. 3b). Interestingly, similar correlations between protein production phases and growth phases were observed in the non-encapsulated bacterial cultures **(Fig. S1a, S1b)**. This indicates that these are the features of the promoter (P_tlpA_) in each strain, which is conserved even when the bacteria are encapsulated and exhibit slower growth and protein production rates.

**Figure 3:**
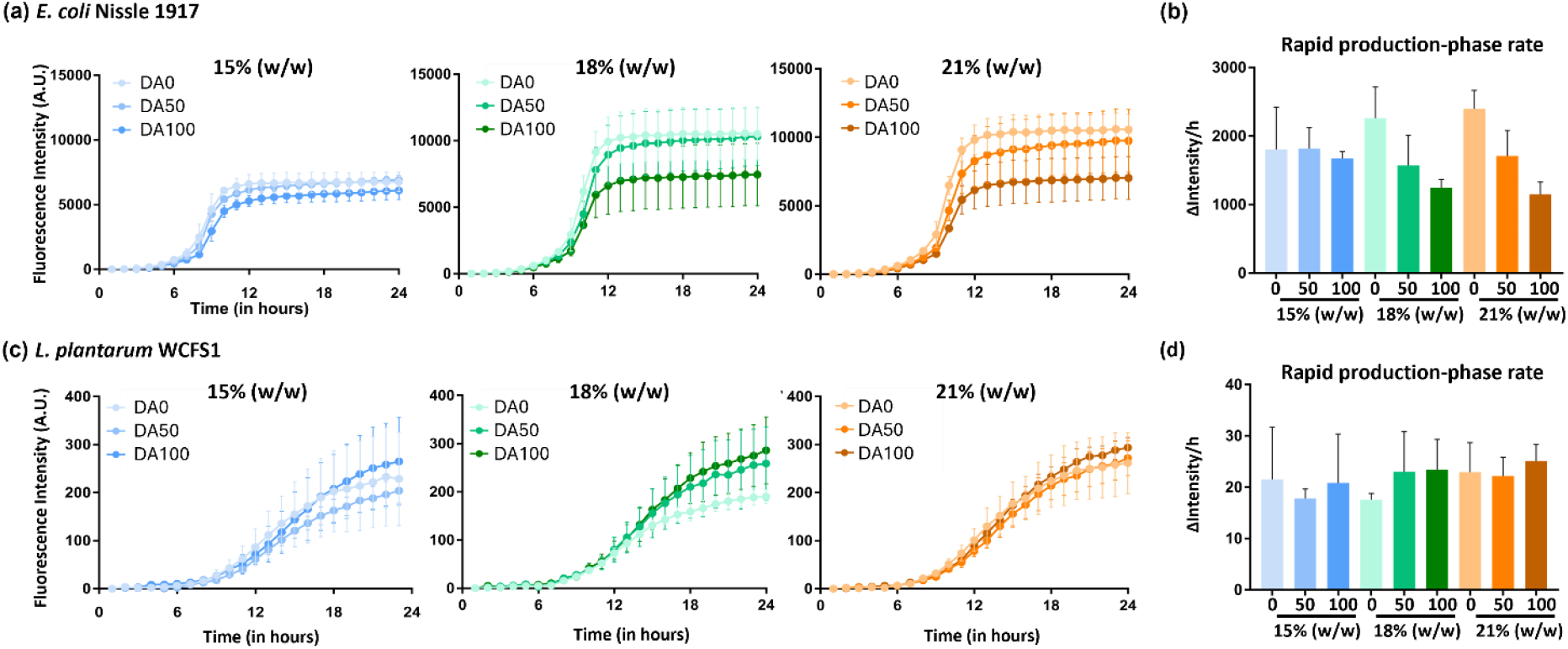
**(a, c)** Normalized kinetics of fluorescence intensity increase in E. coli Nissle 1917 (a) and L. plantarum WCFS1 (c) expressing mCherry in the nine Plu/PluDA combinations for 24 hours (N = 4). **(b**,**d)** Column plots depicting the fluorescence intensity increase rate (change in fluorescence intensity per hour) of E. coli Nissle 1917 (b) and L. plantarum WCFS1 (d) expressing mCherry in the nine Plu/PluDA combinations between the 8-10 hours for E. coli and 10-16 hours for L. plantarum (N = 4).

With *E. coli*, for each polymer concentration, overall protein production and rapid production rate seemed to reduce with increasing degrees of chemical cross-linking especially for 18% and 21% (w/w) formulations **(Fig. 3a)**. In contrast, no apparent trend was observed with *L. plantarum*. Thus, like with growth behavior, mechanical properties of the hydrogels seemed to influence protein production in *E. coli* but not *L. plantarum*.

### 2) Assessment of growth and protein production in bacterial colonies

Within hydrogels, the bacteria are initially homogeneously distributed as single cells, which grow into distinct colonies. After 24 hours of growth, confocal microscopy was used to assess the sizes, morphology, and mCherry production levels of individual colonies. Notably, many more *E. coli* colonies were identified (900 – 1200) compared to *L. plantarum* (140 – 330) **(Table S1)**, suggesting improved adaptation of individual *E. coli* cells to grow by overcoming mechanical restrictions in the hydrogels. In terms of colony volumes, a wide range was observed in all hydrogel formulations, spanning two to three orders of magnitude. With *E. coli*, in all polymer concentrations, the distribution was the largest in DA0 and considerably smaller in DA50 and DA100, as has been observed in our previous study. ^**[9]**^ While these distributions largely overlapped, significant differences of the means were found using the Mann-Whitney test, indicating a trend of decreasing colony volumes on average with higher degrees of chemical cross-linking **(Fig. 4b)**. On the other hand, for *L. plantarum*, the distribution of sizes did not seem to be influenced by the different degrees of chemical cross-linking and a significant increase in mean colony volumes with increasing chemical cross-linking was seen in the 15% and 18% (w/w) polymer concentrations. While this is counter-intuitive, an explanation could be that with greater mechanical restriction, fewer cells grew into colonies and those that did were able to form large colonies on average. This is supported by the fact that the number of colonies in *L. plantarum* that could be measured decreased with increasing chemical cross-linking **(Table S1)**.

**Figure 4:**
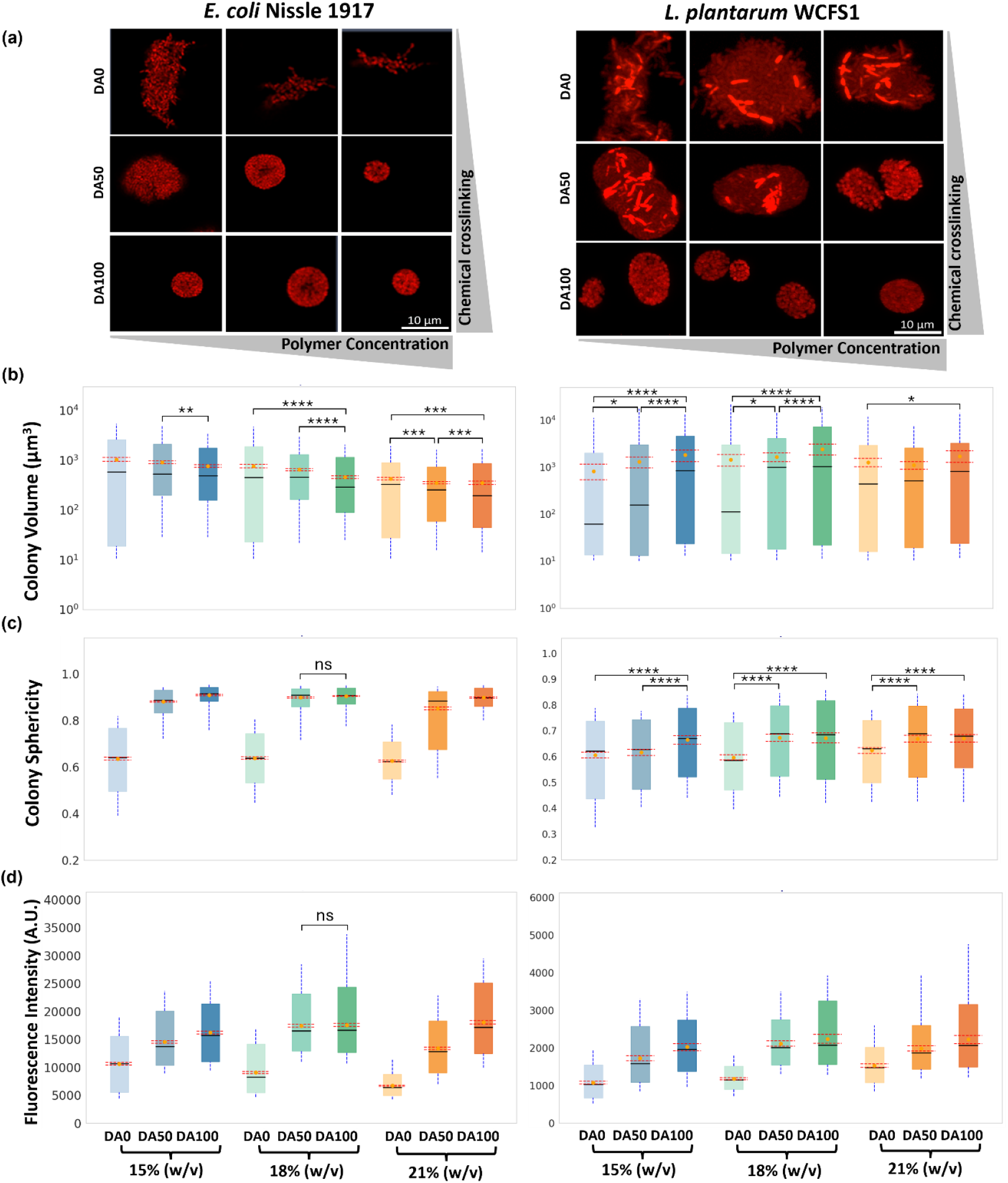
**(a)** Confocal images of representative E.coli Nissle 1917 and L. plantarum WCFS1 colonies in the nine Plu/PluDA combinations after 24 hours of growth. **(b, c, d)** Box plots representing distribution of bacterial colony volume in µm^3^ **(b)**, sphericity **(c)**, and fluorescence intensity **(d)** after 24 hours of growth in the different hydrogels. The graphs on the left represent E.coli Nissle 1917 and the ones on the right are L. plantarum WCFS1. The data was obtained from 4 individual samples and distinct colonies above 10 µm^3^ were quantified. The Mann-Whitney U test was performed to determine significant differences (*p<0.05, **p<0.005, ***p<0.0005, ****p<0.00005). In (c)-E. coli and (d)-both graphs, all distributions within each polymer concentration set were found to be significantly different (p<0.0005), so only non-significant (ns) samples are marked. Comparisons across polymer concentration sets are not indicated.

Apart from affecting growth, mechanical restrictions within the hydrogels were expected to influence colony morphology, so we quantified colony sphericity on a scale of 0 to 1 **(Fig. 4c)**. In DA0 hydrogels with the least mechanical restrictions, both strains exhibited branch-like growth resulting in elongated colonies with low sphericity values (0.6 – 0.7). With *E. coli*, inclusion of chemical cross-linking caused the sphericity of the colonies to drastically increase with values >0.9. In contrast, with *L. plantarum*, the colonies were less branched but remained elongated in DA50 and DA100, resulting in only a slight increase in sphericity. This indicates that in *L. plantarum*, there may be forces exerted between the cells within a colony capable of altering its geometry. This could be a consequence of intercellular interactions in *L. plantarum* that lead to auto-aggregation mediated by adhesive proteins like Aggregation Promoting Factor-D1 (APF-D1).^**[37]**^ While such auto-aggregation has also been reported in *E. coli* Nissle 1917,^**[38]**^ it could be that this phenotype is not manifested under encapsulated conditions or the intercellular forces are not strong enough to alter the geometry of the colony.

Next, we assessed mCherry expression within the colonies to gain insights into the metabolic activity of the encapsulated bacteria **(Fig. 4d)**. The first noticeable difference was that, with *E. coli*, the cells in a colony were relatively homogeneously fluorescent while, with *L. plantarum*, there was considerable heterogeneity. Such population-level differences in fluorescence profiles were also observed in the non-encapsulated bacterial cultures **(Fig. 1d)**, indicating that encapsulation did not alter this property. The next striking observation was that in both strains, the mean fluorescence intensity of the colonies increased with higher degrees of chemical cross-linking. As described in a previous study of ours, ^**[9]**^ this could be due to the mechanically restricted growth allowing the bacteria to dedicate more of its metabolic activity towards the production of the recombinant protein. However, this explanation may not completely justify the effect in *L. plantarum*, since colony volumes are larger in DA50 and DA100. Rather, the growth of fewer colonies with higher degrees of chemical cross-linking may be a stronger contributing factor since the amount of nutrients per colony would be higher, enabling higher levels of recombinant protein production.

All together, these analyses reveal that chemical cross-linking in these hydrogels more strongly influence colony growth in *E. coli* than *L. plantarum*, whereas recombinant protein production is similarly affected in both strains.

### 3) Analysis of metabolic activity by isothermal microcalorimetric analysis

To further investigate the impact of various hydrogel formulations on bacterial metabolism, we employed isothermal microcalorimetry (IMC) using a calScreener device (Symcel, Sweden). This technology enables real-time quantification of metabolic heat generated in the bacterial hydrogels. For this, the bacterial hydrogels were formed in transparent plastic wells and placed in titanium vials, which were arranged in a 48-well format. For each strain, this allowed simultaneous assessment of all hydrogel formulations, non-encapsulated bacterial cultures, and blank medium in duplicates, along with two empty reference wells in each column. The titanium vials were set to maintain the temperature at precisely 37 °C, so metabolic heat was quantified as the heat removed to maintain this temperature. The resulting thermograms show real-time heat flow for 24 hours, in which the total heat is the area under the curve. It is immediately apparent that the metabolic heat generated by *E. coli* is considerably higher than that of *L. plantarum* by a factor of 1.5 – 3 **(Fig. 5a)**. This indicates a higher metabolic rate in the *E. coli* population compared to *L. plantarum* and agrees with the results from Figure 2 showing faster growth of *E. coli* and Figure 3, in which more *E. coli* colonies were identified compared to *L. plantarum*. It also suggests possibly more efficient and versatile use of the available nutrients by *E. coli* for energy production. Moreover, in *E. coli*, chemical cross-linking seems to influence total heat generation, whereas this does not seem to occur in *L. plantarum* **(Fig. 5b)**. For *E. coli*, in 18% and 21% (w/w) gels, those with chemical cross-links (DA50, DA100) exhibit a significant drop in total heat compared to DA0, most likely due to more mechanically restricted growth. In the case of 15% (w/w), the lower total heat values could be attributed to the bacteria requiring less energy to overcome mechanical restrictions for growth. This is supported by the fact that the total heat of non-encapsulated *E. coli* growing in liquid culture is lower than or similar to that of the 15% (w/w) samples **(Fig. S4c)**. This suggests that at higher polymer concentrations, metabolic activity can be influenced by the extent of chemical crosslinking. In *L. plantarum*, the lack of significant differences across all hydrogel formulations aligns with the results in the previous sections indicating minimal influence of the mechanical properties of the hydrogels on their growth. In agreement with this, non-encapsulated *L. plantarum* cultures also generate similar amounts of total heat compared to their encapsulated counterparts **(Fig. S4c)**.

**Figure 5:**
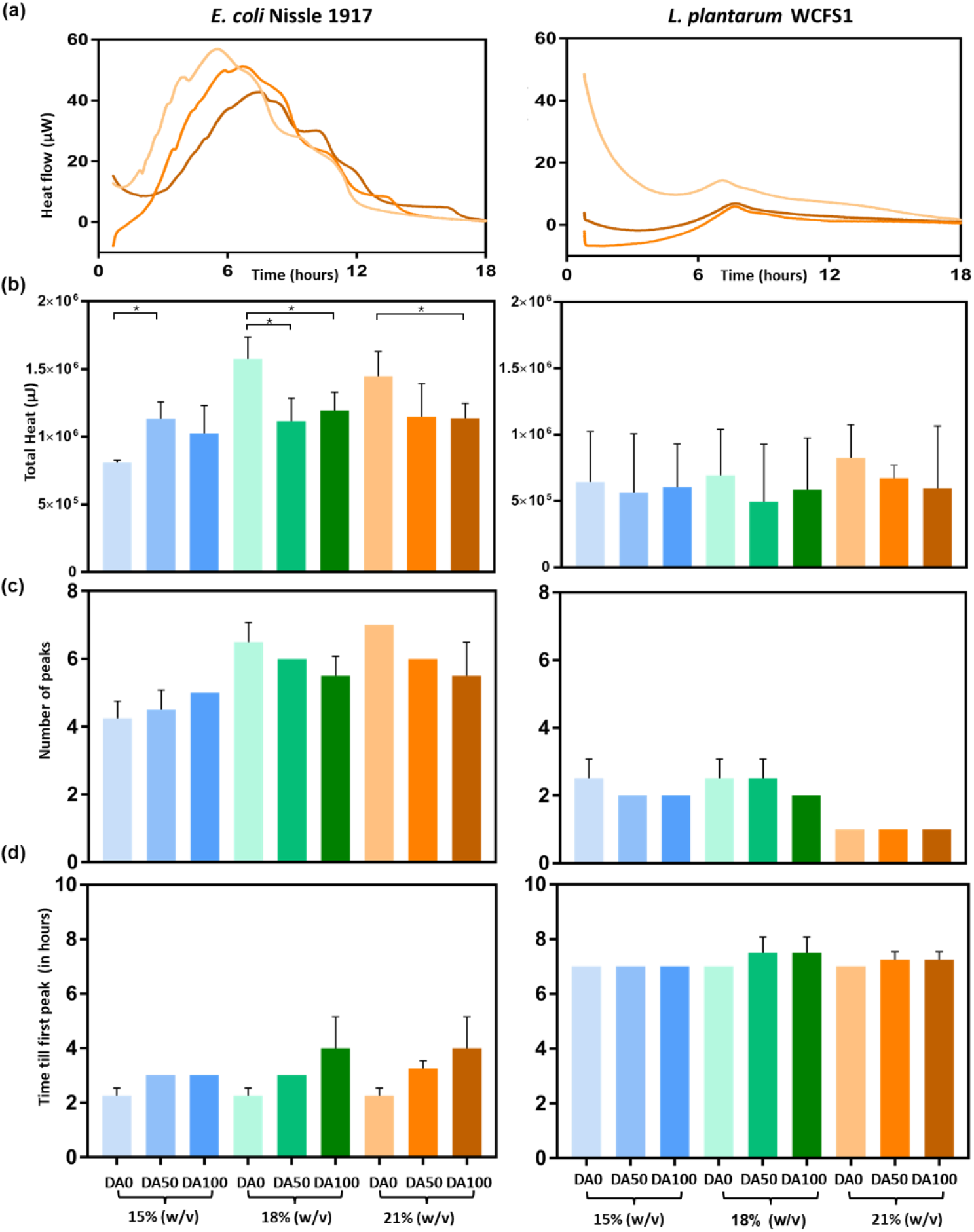
**(a)** Thermograms representing the heat flow (µW) over time from the encapsulated bacteria within 21% w/w Plu/PluDA hydrogels. **(b-d)** Column plots representing total heat (µJ) **(b)**, number of peals **(c)**, and time till first peak **(d)** of each bacterial strain encapsulated in Plu/PluDA gels incubated under isothermal conditions in the CalScreener device at 37 °C for 24 hours. The graph on the left represents E. coli Nissle 1917 and to the right are L. plantarum WCFS1 graphs (N = 4). Analysis of significant differences in the (b) E. coli graph was done using the Mann Whitney U test and have been indicated only for comparisons between different samples within each polymer concentration dataset (*p<0.05). In (b) L. plantarum, the same test revealed non-significant differences between all samples. This analysis was not performed for (c) and (d) due to the semi-quantitative nature of their values.

Apart from total heat, the thermograms also show multiple local maxima, referred to as peaks, that indicate possible transitions in bacterial metabolism (e.g., expansion phase to saturation phase, aerobic to anaerobic, etc.). Once again, there is a clear difference between the two strains in this regard, as *E. coli* exhibits 4 – 7 peaks while *L. plantarum* has 1 – 3 **(Fig. 5c)**. This suggests that *E. coli* is able to adapt to the mechanical restrictions and, eventually, nutrition limitations by switching between different metabolic pathways, whereas *L. plantarum* does not have this versatility. In *E. coli*, this is further supported by the fact that more peaks are observed in 18% and 21% (w/w) hydrogels. However, chemical cross-linking in these hydrogels reduce the number of peaks by at least one, suggesting that a mechanical barrier is probably reached beyond which further growth cannot be achieved through metabolic changes. The inference that *E. coli* is adapting to overcome mechanical restrictions is further supported by the fact that the total heat detected from hydrogels **(Fig. 5b)** is double the values detected in non-encapsulated liquid cultures **(Fig. S4c)**. Furthermore, more peaks are seen before the global maxima with the encapsulated *E. coli* (**Fig. 5a**) compared to those that were not encapsulated (**Fig. S4a**). In contrast, with *L. plantarum*, the total heat and peak occurrence are similar in both hydrogels and non-encapsulated liquid cultures. In all cases, the time till the first peak represents the duration till the first metabolic transition occurs. In *E. coli*, this occurs as soon as 2 hours in the mechanically less restricted hydrogels and increases to 4 hours with higher degrees of chemical cross-linking in the 18% and 21% (w/w) hydrogels **(Fig. 5d)**. In non-encapsulated liquid cultures, the time to first peak is 2 hours for *E. coli* **(Fig. S4a)**. This suggests that *E. coli* is possibly adapting to nutrient limitations sooner than mechanical restrictions in the 15% (w/w) and DA0 hydrogels, while slower growth in the more mechanically restricting hydrogels possibly delays the need for this adaptation. In *L. plantarum*, time to first peak occurs almost uniformly around 7 - 8 hours in all hydrogel formulations **(Fig. 5d)** and non-encapsulated cultures **(Fig. S4b)**, which aligns with the observation throughout this study that mechanical restrictions within these hydrogels do not considerably influence growth and metabolic activity in this strain. The only parameter that chemical cross-linking seems to considerably affect in *L. plantarum* seems to be recombinant protein production levels in colonies as observed in Fig. 4d, but IMC analysis reveals that this is not due to any major shift in overall metabolic activity. Therefore, the difference is possibly only due to an efficient realignment of resources towards recombinant protein production when colony growth or numbers are lower.

## Conclusions

This study provides a comprehensive comparison of the adaptive responses of probiotic gram-negative *E. coli* Nissle 1917 and gram-positive *L. plantarum* WCFS1 to mechanical confinement within hydrogel matrices. Our findings reveal that while both bacterial species can grow and produce recombinant proteins under confinement, their responses to the mechanical properties of the encapsulating material differ significantly. Time-resolved analysis of growth and recombinant protein production showed that encapsulation slows down and delays the different phases in both bacteria, with *L. plantarum* being natively slower in both parameters. However, the correlations between protein production and growth phases seem to be conserved, revealing that encapsulation does not alter the performance of the promoter in both bacteria. Through confocal microscopy analysis, we observed that *E. coli* exhibited notable changes in colony growth, size, morphology, and metabolic activity in response to varying hydrogel stiffness, indicating a greater sensitivity to mechanical constraints. In contrast, *L. plantarum* showed a more robust growth pattern with minimal changes in colony morphology and metabolic activity, likely due to its thicker cell wall and slower growth rate, which may confer an advantage in overcoming mechanical restrictions. However, colonies of both bacteria exhibited increased recombinant protein production with higher matrix stiffness, which underscores the potential of optimizing hydrogel properties to enhance the functionality of ELMs. These insights into species-specific adaptations to mechanical confinement advance our understanding of the interplay between material properties and microbial physiology. This knowledge is crucial for the rational design of ELMs tailored for therapeutic applications, where precise control over the behavior of engineered probiotic bacteria is essential. Overall, this study highlights the importance of considering the mechanical environment in the development of ELMs and provides a foundation for future research aimed at optimizing material-microbe interactions for various applications.

## Acknowledgements

We thank Farrukh Usama and Silke Siegrist from the group of Prof. Aránzazu del Campo for generously providing Pluronic F127 diacrylate. We thank Dr. Shardul Bhusari for his support with rheological experiments. We also thank Dr. Kasper Kragh and Moa Persson from Symcel AB for their generous support regarding calScreener data. Biorender was used for generating the schemes in this manuscript.

## Data Availability

The information provided in the main text and supporting information are sufficient to reproduce the experiments described in this study. The raw and processed datasets along with relevant metadata are available from the corresponding author upon reasonable request.

