## Supplementary information for "Adaptations of gram-negative and gram-positive probiotic bacteria in engineered living materials"

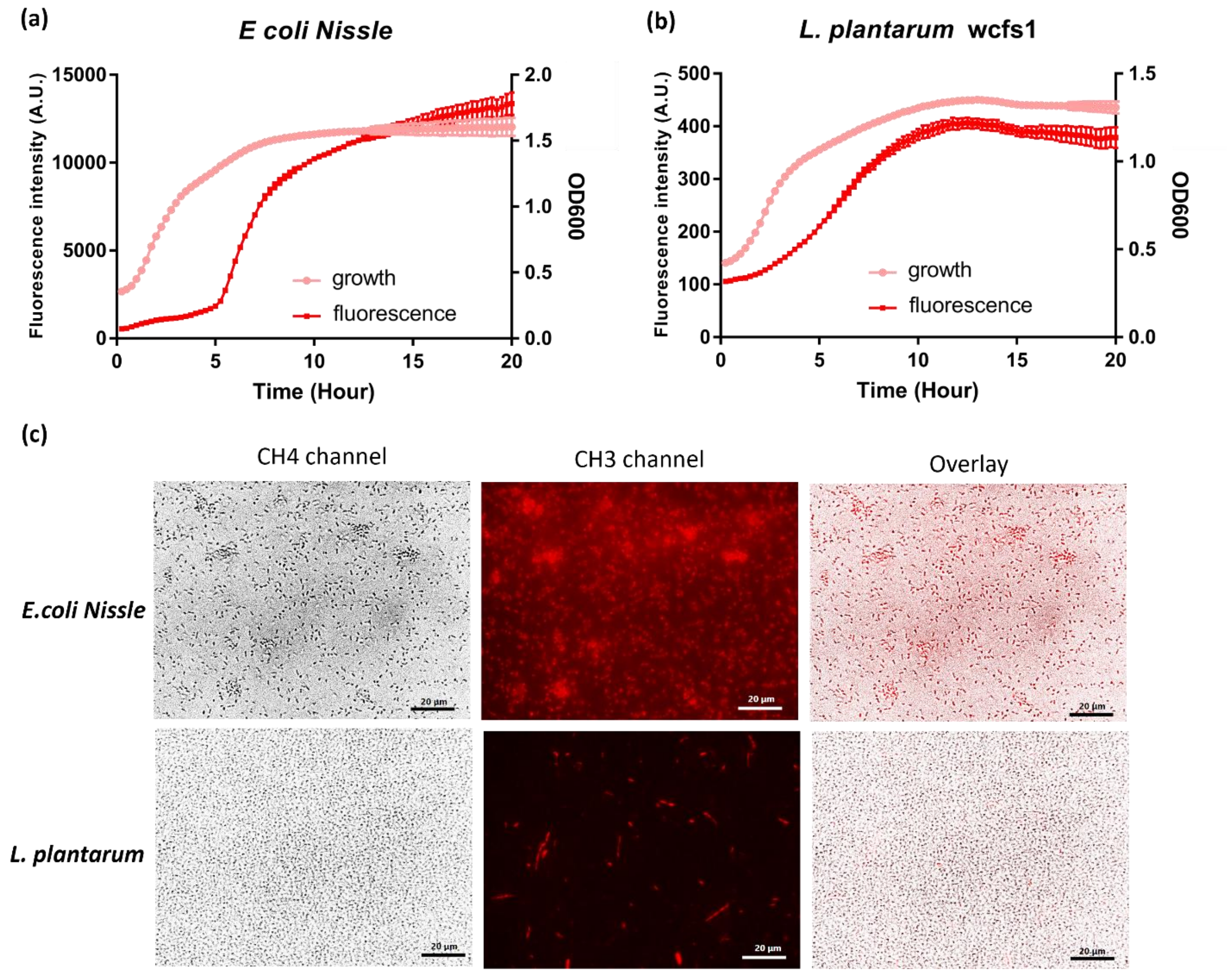


***Figure S1:*** *Plots showing the growth kinetics and fluorescence expression kinetics of (a) E. coli Nissle 1917 expressing mCherry, and (b) L. plantarum WCFS1 expressing mCherry. (c) Microscopy images from cultures of E. coli and L. plantarum expressing mCherry after 20 hours of growth kinetics.*

| Poly. conc. (w/w) | Chem. cross-link. | *E. coli* | *L. plantarum* |
|---|---|---|---|
| **15%** | DA0 | 1014 | 333 |
| | DA50 | 1249 | 307 |
| | DA100 | 1174 | 143 |
| **18%** | DA0 | 1102 | 314 |
| | DA50 | 992 | 203 |
| | DA100 | 1033 | 142 |
| **21%** | DA0 | 1237 | 246 |
| | DA50 | 904 | 236 |
| | DA100 | 832 | 169 |

***Table S1:*** *List of number of colonies analysed per condition of E. coli and L. plantarum to determine bacterial colony volume, colony sphericity and fluorescence intensity mean.*

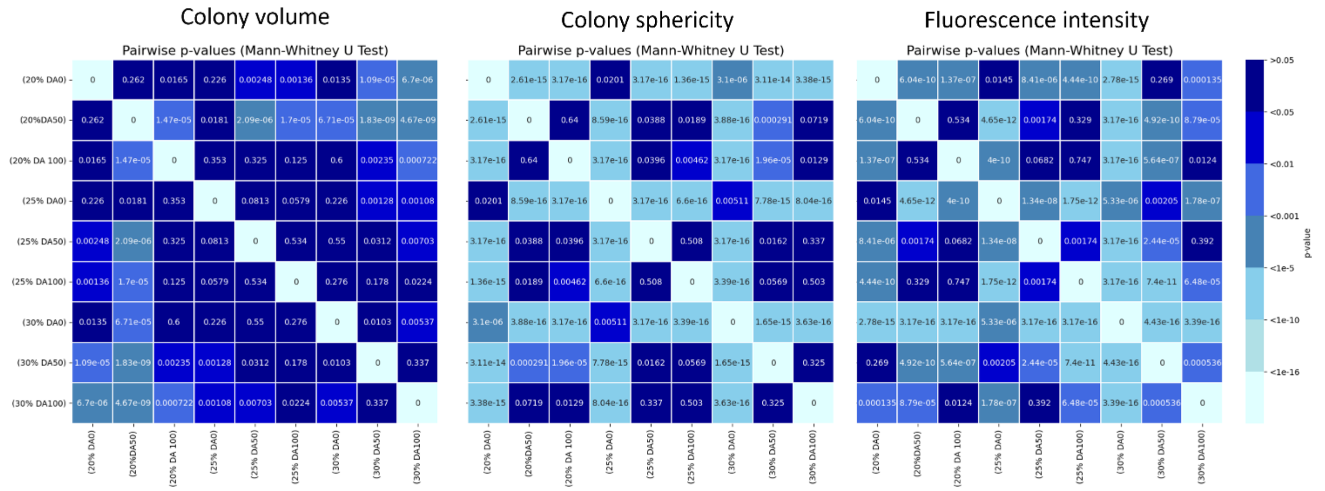


***Figure S2:*** *Heatmaps representing the statistical significance in terms of P-values obtained using Mann-Whitney U test, to assess pairwise differences between nine different formulations for each of the three parameters assessed for E.coli Nissle 1917: Colony volume, Colony sphericity and Fluorescence intensity.*

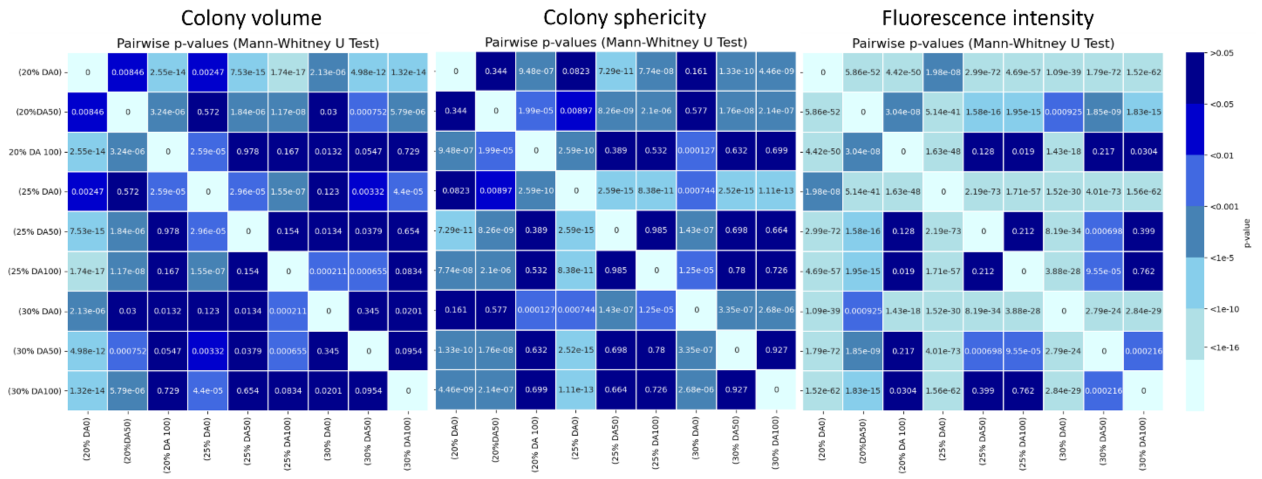


***Figure S3:*** *Heatmaps representing the statistical significance in terms of P-values obtained using Mann-Whitney U test, to assess pairwise differences between nine different formulations for each of the three parameters assessed for L. plantarum WCFS1: Colony volume, Colony sphericity and Fluorescence intensity.*

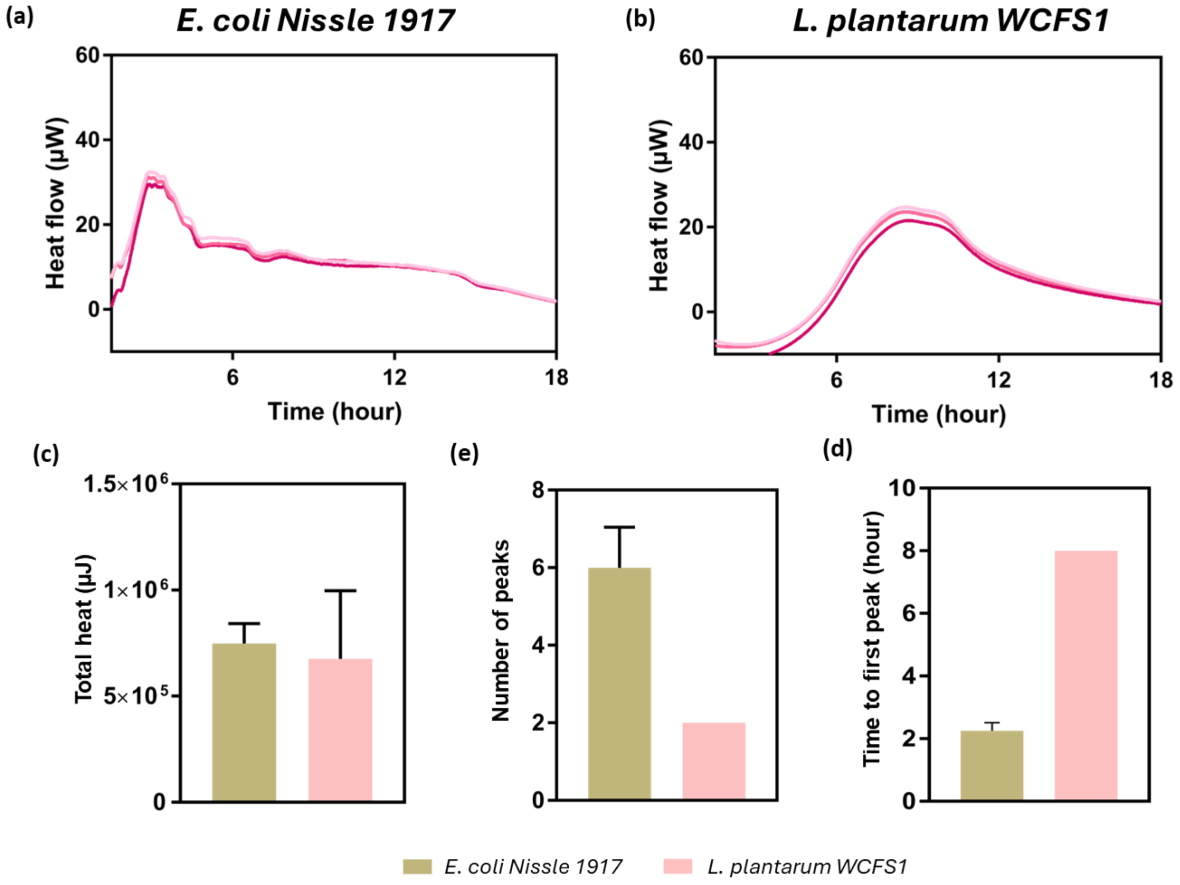


***Figure S4: (a,b)*** *Thermograms depicting heat flow quantified from liquid cultures of E. coli Nissle 1917 (a) and L. plantarum WCFS1 (b), using calScreener.* ***(c,d,e)*** *Column plots representing total heat (c), number of peaks (d), and time to first peak (e), determined from the thermograms.*
